## Supplementary figures and images for "Drug repurposing for Cystic Fibrosis: identification of drugs that induce CFTR-independent fluid secretion in nasal organoids"

### Supplemental figures

#### Supplemental figure S1

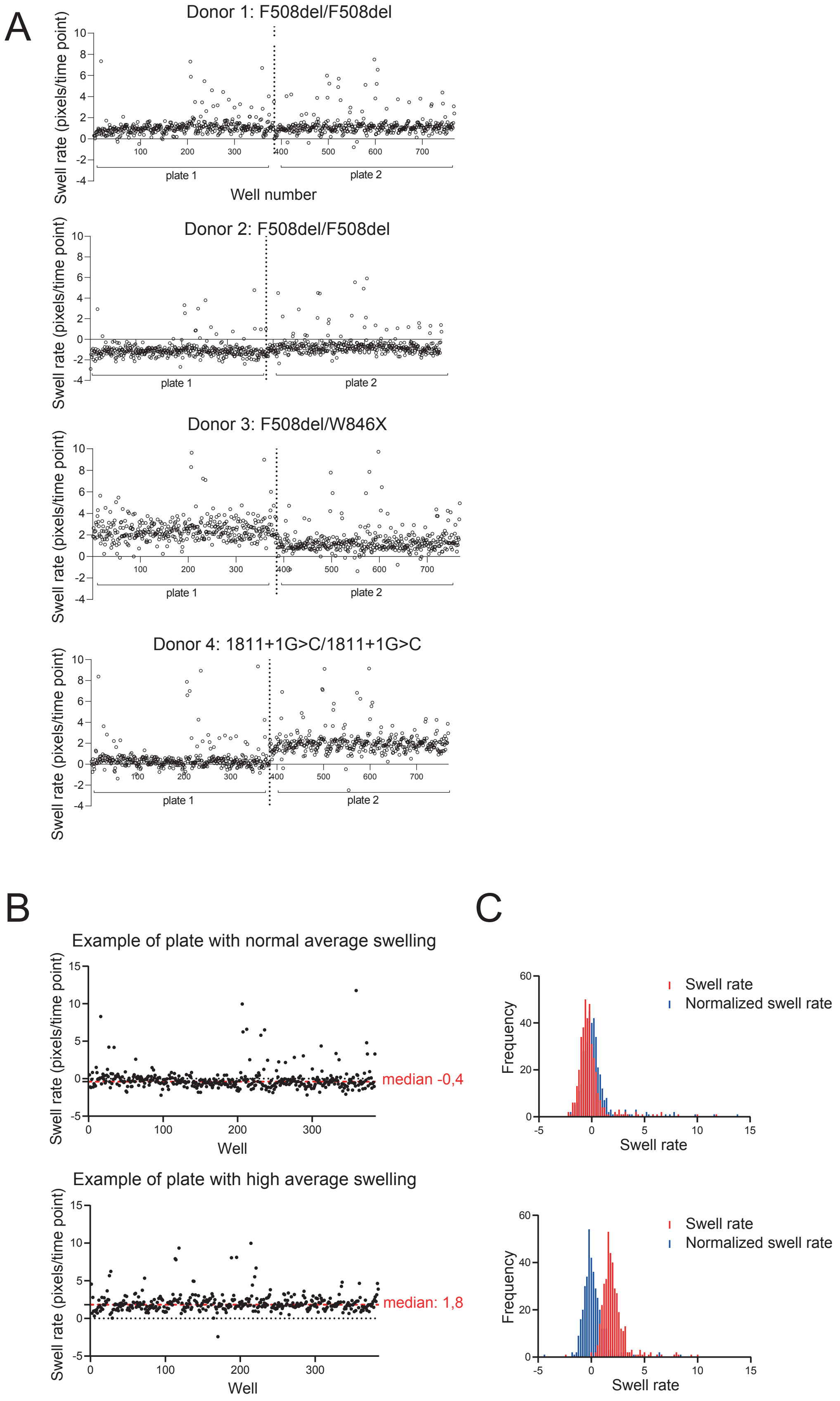

#### Supplemental figure S2

A

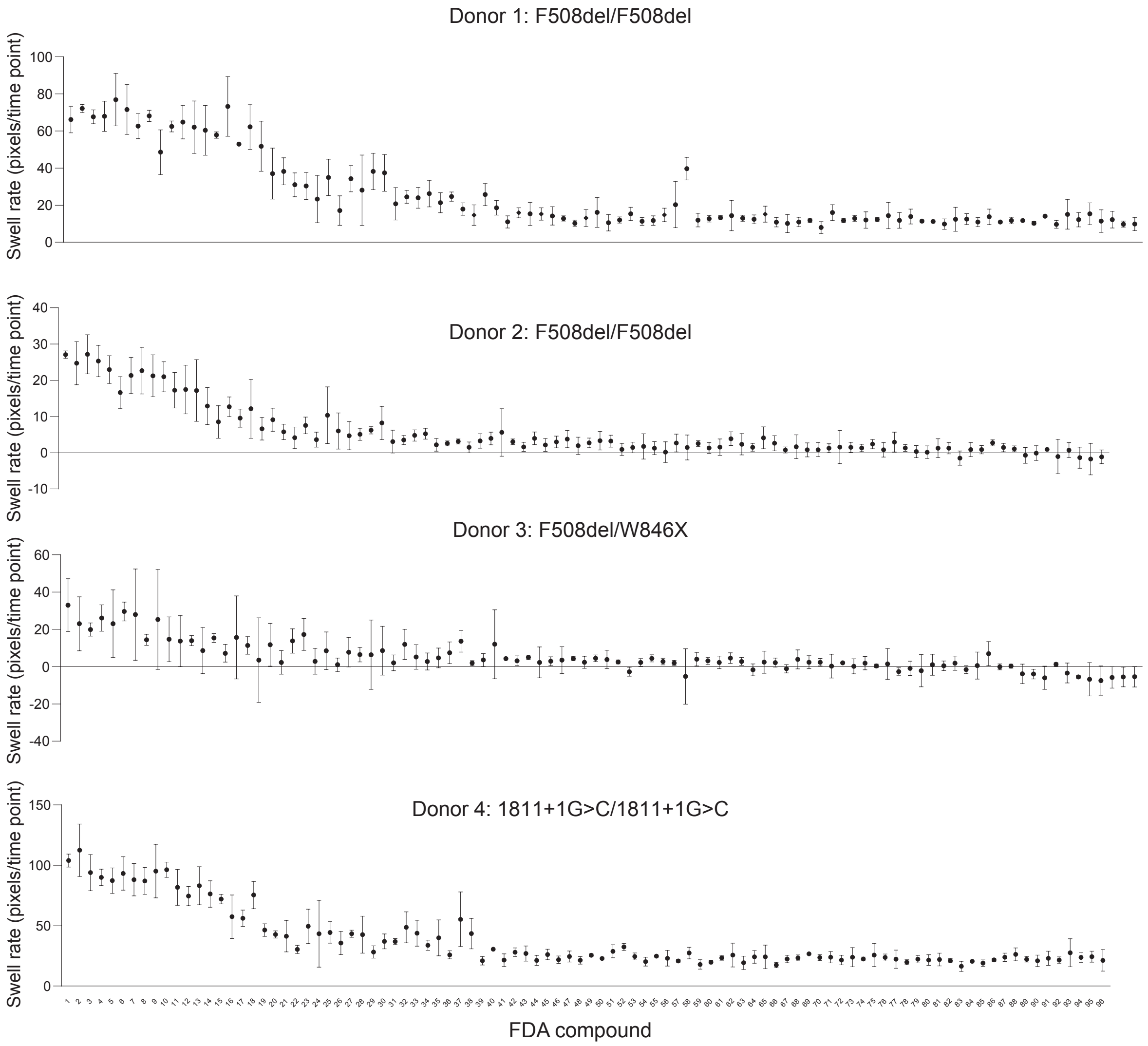

#### Supplemental figure S3

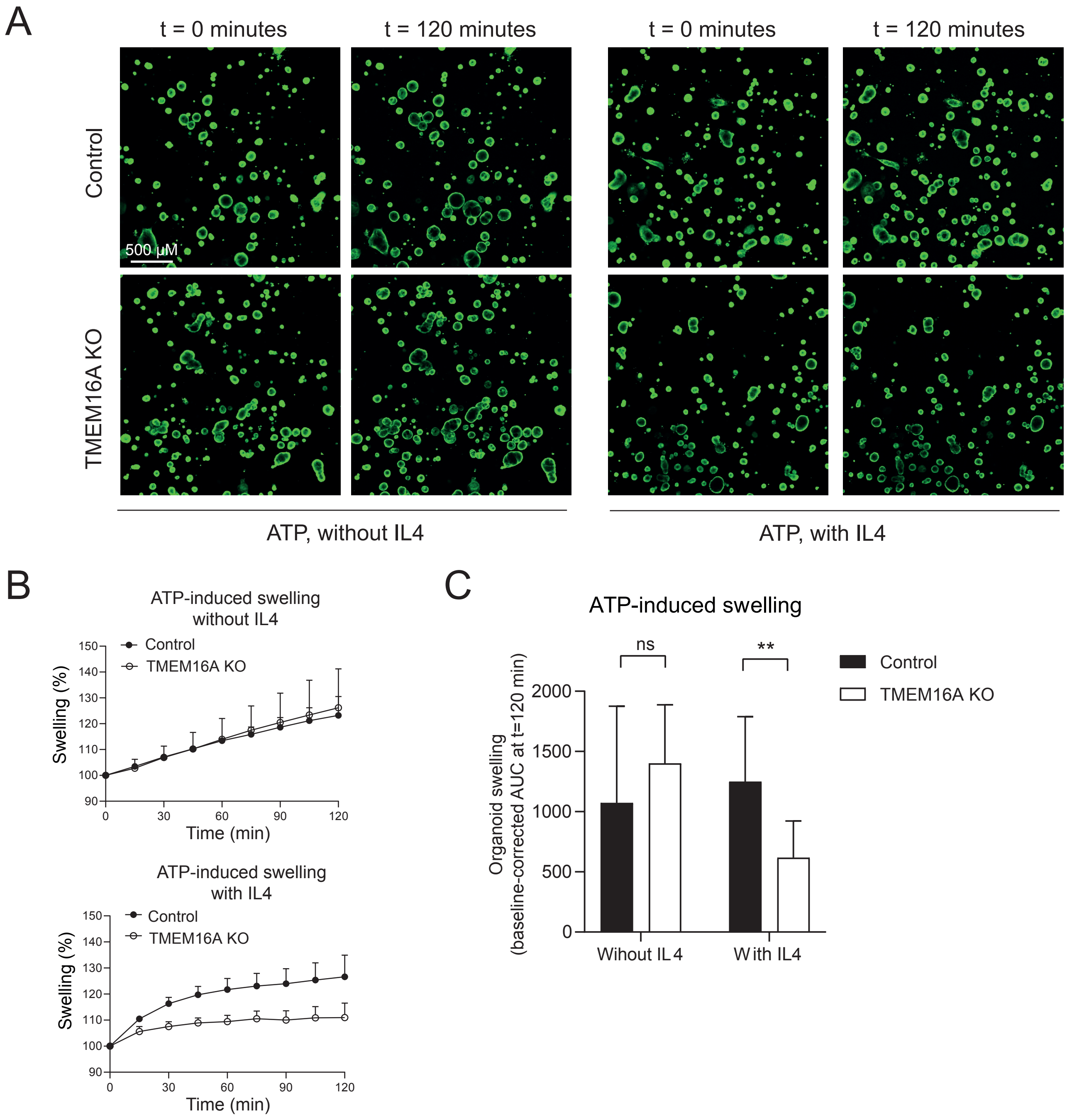

#### Supplemental figure S4

A

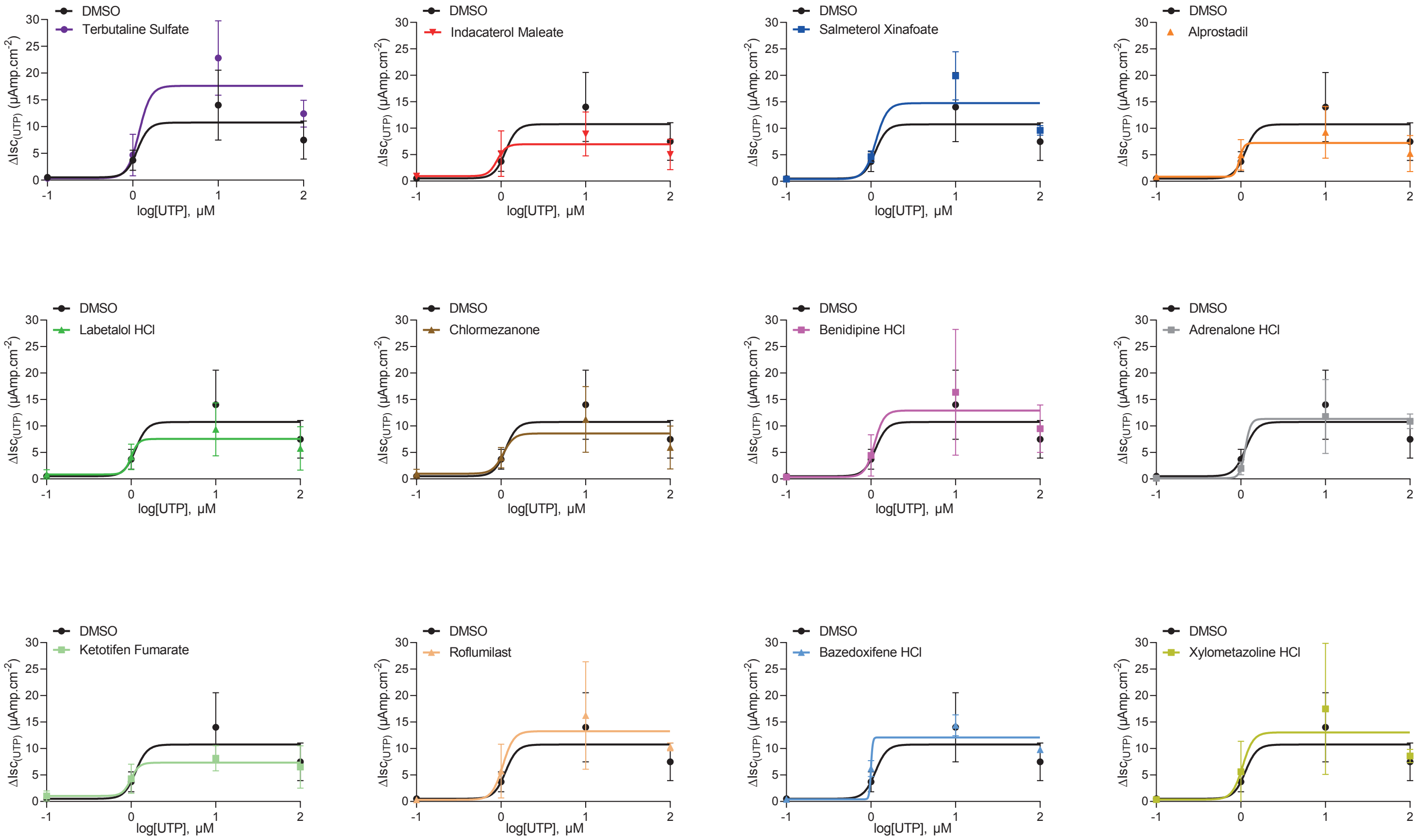


B

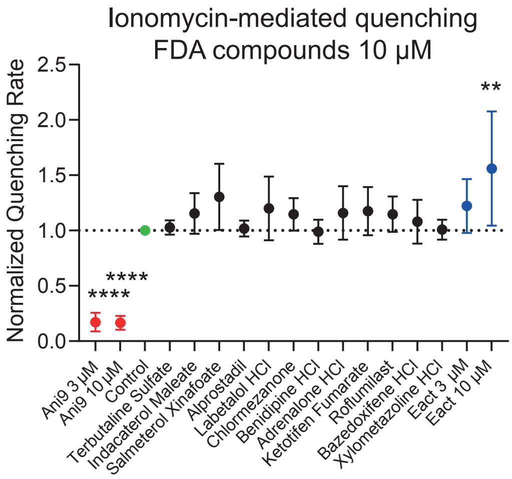
