## Supplemental tables for "Drug repurposing for Cystic Fibrosis: identification of drugs that induce CFTR-independent fluid secretion in nasal organoids"

**Table S1.** Basal cell isolation and expansion medium

| Reagents | Concentration | Company |
| --- | --- | --- |
| Bronchial epithelial cell medium-basal (BEpiCM-b) | 50 % (v/v) | ScienCell, Carlsbad, CA, USA |
| Advanced DMEM/F12 | 23.5 % (v/v) | Gibco, Waltham, MA, USA |
| B-27 Supplement, serum free | 2 % (v/v) | Gibco, Waltham, MA, USA |
| GlutaMAX Supplement | 1 % (v/v) | Gibco, Waltham, MA, USA |
| HEPES (1 M) | 10 mM | Gibco, Waltham, MA, USA |
| (±)-Epinephrine hydrochloride | 0.5 μg/mL | Sigma-Aldrich, St Louis, MO, USA |
| Hydrocortisone | 0.5 μg/mL | Sigma-Aldrich, St Louis, MO, USA |
| 3,3′,5-Triiodo-L-thyronine sodium salt | 100 nM | Sigma-Aldrich, St Louis, MO, USA |
| N-Acetyl-L-cysteine | 1.25 mM | Sigma-Aldrich, St Louis, MO, USA |
| Nicotinamide | 5 mM | Sigma-Aldrich, St Louis, MO, USA |
| SB 202190 (p38i) | 500 nM | Sigma-Aldrich, St Louis, MO, USA |
| DMH-1 (BMPi) | 1 μM | Selleck chemicals, Planegg, Germany |
| A83-01 (TGF-βi) | 1 μM | Tocris, Bristol, UK |
| Y-27632 (ROCKi) | 5 μM | Selleck chemicals, Planegg, Germany |
| DAPT (NOTCHi; only in expansion medium) | 5 μg/mL | Fisher Scientific, Landsmeer, Netherlands |
| Recombinant human FGF-7 | 25 ng/mL | Peprotech, Rocky Hill, NJ, USA |
| Recombinant human FGF-10 | 100 ng/mL | Peprotech, Rocky Hill, NJ, USA |
| Recombinant human EGF | 5 ng/mL | Peprotech, Rocky Hill, NJ, USA |
| Recombinant human HGF | 25 ng/mL | Peprotech, Rocky Hill, NJ, USA |
| Rspondin 1 conditioned medium (from Rspo1 cells Cultrex®) | 20 % (v/v) | Trevigen, Gaithersburg, MD, USA |
| Penicillin-Streptomycin | 1 % (v/v) | Gibco, Waltham, MA, USA |
| Primocin | 100 μg/mL | Invivogen, San Diego, CA, USA |
| Amphotericin B (only in isolation medium) | 250 μg/mL | Gibco, Waltham, MA, USA |
| Gentamicin (only in isolation medium) | 50 μg/mL | Sigma-Aldrich, St Louis, MO, USA |
| Vancomycin (only in isolation medium) | 50 μg/mL | Sigma-Aldrich, St Louis, MO, USA |

**Table S2.** ALI differentiation medium

| Reagent | Concentration | Company |
| --- | --- | --- |
| Advanced DMEM/F12 | 98.5% (v/v) | Gibco, Waltham, MA, USA |
| (±)-Epinephrine hydrochloride | 0.5 μg/mL | Sigma-Aldrich, St Louis, MO, USA |
| Hydrocortisone | 0.5 μg/mL | Sigma-Aldrich, St Louis, MO, USA |
| 3,3′,5-Triiodo-L-thyronine sodium salt | 100 nM | Sigma-Aldrich, St Louis, MO, USA |
| Penicillin-Streptomycin | 1 % (v/v) | Gibco, Waltham, MA, USA |
| A83-01 (TGF-βi) | 50 nM | Tocris, Bristol, UK |
| TTNPB (Retinoic acid agonist) | 100 nM | Cayman, Ann Arbor, MI |
| Recombinant human EGF | 0.5 μg/mL | Peprotech, Rocky Hill, NJ, USA |

**Table S3.** Airway organoid medium

| Reagent | Concentration | Company |
| --- | --- | --- |
| Advanced DMEM/F12 | 95.5% (v/v) | Gibco, Waltham, MA, USA |
| B-27 Supplement, serum free | 2 % (v/v) | Gibco, Waltham, MA, USA |
| GlutaMAX Supplement | 1 % (v/v) | Gibco, Waltham, MA, USA |
| HEPES | 10 mM | Gibco, Waltham, MA, USA |
| N-Acetyl-L-cysteine | 1.25 mM | Sigma-Aldrich, St Louis, MO, USA |
| Nicotinamide | 5 mM | Sigma-Aldrich, St Louis, MO, USA |
| SB 202190 (p38i) | 500 nM | Sigma-Aldrich, St Louis, MO, USA |
| A83-01 (TGF-βi) | 500 nM | Tocris, Bristol, UK |
| Y-27632 (ROCKi) | 5 μM | Selleck chemicals, Planegg, Germany |
| Penicillin-Streptomycin | 1 % (v/v) | Gibco, Waltham, MA, USA |
| Recombinant human FGF-7 | 5 ng/mL | Peprotech, Rocky Hill, NJ, USA |
| Recombinant human FGF-10 | 10 ng/mL | Peprotech, Rocky Hill, NJ, USA |

**Table S4.** Antibodies

| Antibody | Source | Identifier | Dilution |
| --- | --- | --- | --- |
| Mouse anti-MUC5AC | Thermo Fischer Scientific | #MA1-38223 | 1:500 |
| Rabbit anti-β-tubulin IV | Abcam, Cambridge, UK | #ab179509 | 1:500 |
| Rabbit anti-TMEM16A | Abcam, Cambridge, UK | AB64085 | 1:500 |
| Rabbit anti-HSP90 | Developed by laboratory of Prof. I. Braakman | NA | 1:10.000 |
| Goat anti-mouse IgG1, Alexa Fluor 647 | Invitrogen, Waltham, MA, USA | A-21240 | 1:500 |
| Goat anti-rabbit IgG, Alexa Fluor 488 | Invitrogen, Waltham, MA, USA | A-11034 | 1:500 |
| Goat anti-rabbit Immunoglobulins/HRP | Dako, Santa Clara, CA, USA | P0448;RRID:AB_2617138 | 1:2.000 |

**Table S5.** qPCR primers

| Gene | Forward primer (5’- 3’) | Reverse primer (5’- 3’) |
| --- | --- | --- |
| *ANO1* | AGGATTCCTTTTTCGACAGCAA | CGTTTTCACCGTTGTAGTCTCC |
| *SLC26A9* | GACTACATCATTCCTGACCTGC | AGGAGTAGAGGCCATTGACTG |
| *SLC26A4* | TGGTGGCTTGCAGATTGGAT | AGCTGTGAGACCAGCACTTG |
| *CLCN2* | TTGATCCTGCTCCCTTCCAG | CATAAGCATGGTCCACTCCC |
| *SCNN1A* | TCTGCACCTTTGGCATGATGT | GAAGACGAGCTTGTCCGAG |
| *CFTR* | CAACATCTAGTGAGCAGTCAGG | CCCAGGTAAGGGATGTATTGTG |
| *ATP5B* | TCACCCAGGCTGGTTCAGA | AGTGGCCAGGGTAGGCTGAT |
| *RPL13A* | AAGGTGGTGGTCGTACGCTGTG | CGGGAAGGGTTGGTGTTCATCC |
