## Supplementary material for "Drug repurposing for Cystic Fibrosis: identification of drugs that induce CFTR-independent fluid secretion in nasal organoids": Figure legends of supplemental figures

**Figure S1**. Primary screening assay in individual CF donors. (**a**) The primary screening assay was performed in four CF donors with different CFTR genotypes as indicated in the graphs (n=4 independent donors, 1-3 replicates per donor). Organoid swelling was determined in 384-wells plate format with 2 FDA compounds combined in a single well. Graphs depict the mean swell rate of all organoids in a well. Dotted lines represent different assay plates; (**b**) A plate normalization step was performed as median swell rates varied between assay plates and donors. Examples are shown from a plate with normal median swelling (upper panel) and from a plate with high median swelling (lower panel); (**c**) Histograms show the distribution of measurements before (red) and after (blue) plate normalization.

**Figure S2.** Secondary screening assay in individual CF donors. (**a**) The secondary screening assay was performed in the more conventional 96-wells plate format in the same four CF donors as the primary screening assay, as indicated in the graphs. Organoid swelling is shown for individual donors.

**Figure S3**. ATP-induced swelling in TMEM16A KO nasal organoids. (**a**) Representative confocal images (G542X/CFTRdele2.3(21kb)) of TMEM16A KO and control nasal organoids after stimulation with ATP. Organoids were treated with or without IL-4 for 48h; (**b,c**) Quantification of ATP-induced swelling in TMEM16A KO and control nasal organoids, treated without (upper panel) or with (lower panel) IL-4 for 48h (n=3 independent CFTR-null donors, 1-7 measurements per donor). Analysis of difference was performed using unpaired t-tests. ** p < 0.01.

**Figure S4.** Effect of the hit compounds on TMEM16A in other *in vitro* model systems. (**a**) The effect of hit compounds on UTP-dose response curves was analyzed in Ussing Chamber measurements with CFTR-null nasal cells. Data is shown for incubation with DMSO alone or with one of the 12 hit compounds. Nonlinear regression was used to fit a dose-response curve (n=3 independent donors, each compound was measured in at least 2 different donors; n =2-3 measurements per donor); (**b**) The Ionomycin-induced quenching assay in CFBE cells was also performed with a higher concentration of the FDA hit compounds (10 µM instead of 3 µM). TMEM16A-dependency was demonstrated with sensitivity for Ani9 (3 and 10 uM, shown in red). DMSO was used as negative control (shown in green) and E_act_ (3 and 10 uM) as positive control (shown in blue). For quantification, quenching rates were normalized to the control and a one-way ANOVA with Dunnett’s post-hoc test was used to analyze differences with control. ** p < 0.01, **** p<0.0001.
